# Nutrient niche differentiation among European herbaceous species reflects an extinction–invasion continuum

**DOI:** 10.1101/2025.08.20.670250

**Authors:** Daniil J.P. Scheifes, Harry Olde Venterink, Hugo J. de Boer, Paul M. J. Berghuis, Hans Lambers, Julian Schrader, Mariska te Beest, Karin T. Rebel, Martin J. Wassen

## Abstract

Threatened and invasive plant species may appear worlds apart; however, we propose that mechanisms underlying invasive success and extinction risk among European herbaceous species constitute a continuum from successful, well-dispersing, fast-growing species to threatened, slow-growing species. We provide empirical evidence for such an extinction-invasion continuum and show that threatened and naturalized invasive plant species occur at opposite ends. Threatened species persist in phosphorus-limited nutrient-poor habitats, while naturalized and invasive species more often occur in nitrogen-limited nutrient-rich habitats. These opposing niches suggest invasive species do not directly displace threatened species; instead, species replacement and extinction result from nutrient regime shifts. Mitigating and preventing nutrient enrichment, especially phosphorus, for nature conservation protects existing nutrient niches for threatened species and limits plant invasion.

---

Ecosystem invasions by non-native plant species have risen steeply over the last century and are still accumulating (1) while the number of threatened plant species is also rising due to anthropogenic activities (2). Non-native plant invasion is often mentioned as one of the main driving factors for diversity loss (2) and species extinction (3), but whether they pose a direct threat to threatened species remains debated (4, 5). Several studies have aimed to identify the key functional traits and habitat preferences that can distinguish invasive from threatened species. Considering invasive and naturalized species, studies have indicated that these species tend to invest more in sexual reproduction (6–8) and follow a fast-growth strategy (9–12), making them successful in disturbed and nutrient-rich habitats (13, 14). In contrast, threatened species typically invest less in sexual reproduction and have a slow-growth strategy (15, 16), often persisting in nutrient-poor habitats (14, 17). Furthermore, threatened species particularly persist under phosphorus (P)-limited growth conditions in both Eurasian herbaceous ecosystems (18) and a smaller set of Brazilian cerrado plants (14), as indicated by a high nitrogen (N) to P stoichiometric ratio in the community biomass.

It has been hypothesized that general mechanisms underlying threatened status are similar to those explaining invasive success and are therefore two sides of the same coin (19, 20). Recent studies on plant dispersal and naturalization in Europe suggest that this hypothesis might apply to plants, as successful species in their native range also expand outside their native range (21, 22). However, these studies did not directly compare threatened and invasive species. Naturalized and invasive species appear to differ from threatened species in their nutrient niche and functional strategies, but comparative studies on this topic are rare. While some studies have compared traits between non-invasive and invasive plant species (11) and others have compared nutrient niches and traits between threatened and non-threatened plant species (15), large-scale, integrated comparisons are still lacking. Such comparisons could provide important information for both controlling plant invasion and preventing extinction by identifying shared drivers that can help mitigate the impact of invasive species while enhancing conservation efforts for threatened species.

Here we propose arranging plant species along an extinction–invasion continuum that reflects a gradient of success, ranging from species that decline, to those that are stable within their native range, to species that are successful across the globe (cf. Fig. 1). Such a continuum allows for the comparison of groups of species based on their functional strategies and nutrient requirements, and the evaluation of the extent to which invasive plant species may pose a direct threat to native species.

**Figure 1.**
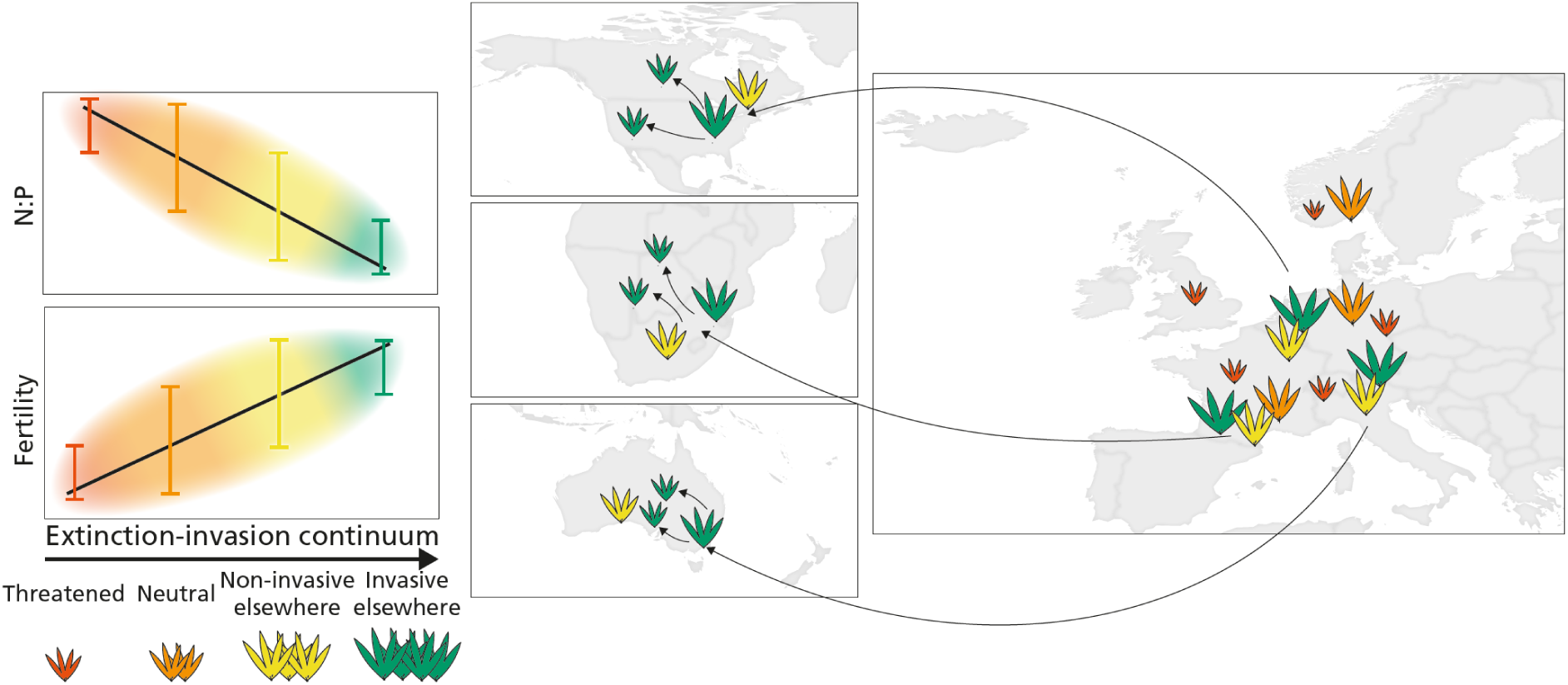
| Conceptual model explaining the extinction-invasion continuum. The continuum orders European species between threatened species (red) that persist but are in decline; neutral species (orange) that are stable in Europe and are neither threatened nor have they naturalized outside Europe; non-invasive species elsewhere (yellow) that have naturalized outside Europe such as North America, South Africa and Australia but are not invasive and invasive species elsewhere (green) that have naturalized outside Europe but have spread over considerable distances. The left panels illustrate our expectation that species along the increasing extinction-invasion continuum have a lower nitrogen to phosphorus stoichiometric ratio (N:P) and occupy more fertile niches.

Using a unique dataset of 837 European sites of herbaceous plant communities, including 682 plant species, we examined species composition, community aboveground biomass and community aboveground N and P concentrations. From this, we determined nutrient availability niches (community biomass and Ellenberg’s fertility indicator (23)) and N:P stoichiometric niches. We retrieved plant traits related to growth rate and life history (building on previous studies (15, 24) with additional trait data). We compared these nutrient niches and traits along species’ positions within an extinction-invasion continuum. Positions along this continuum were approached using four categories: (I) species with a threatened status according to regional red lists, (II) species that are not threatened and have not naturalized outside Europe, (III) species that have naturalized outside Europe but are not considered invasive (25), and (IV) species that have naturalized and are considered invasive outside Europe (26).

We expected two patterns to emerge when comparing species’ nutrient niches and traits of the species along their positions on the extinction-invasion continuum. We expected species to occupy distinct positions along this continuum, first determined by their nutrient availability niches (reflected in community biomass and Ellenberg’s fertility indicator) and their N:P stoichiometry (Fig. 1); and second, by their functional traits related to above- and belowground growth and life history, as species have evolved contrasting strategies in response to infertile, P-limited versus fertile, N-limited conditions (Fig. S1).

## Nutrient niches and traits along the extinction-invasion continuum

Our results demonstrate that (I) nutrient niches and various traits related to growth rate and life history strongly cluster along the proposed extinction-invasion continuum and (II) threatened and invasive species operate within distinct nutrient niches. First, we found that this continuum is linked to variables related to overall site fertility and productivity (Fig. 2a, b) as well as nutrient availability and stoichiometry (Fig. 2c, d, e). Species’ mean community biomass (Fig. 2a, g m^-2^) and Ellenberg’s soil fertility indicator (Fig. 2b) increased along this continuum. Species mean community P concentration (Fig. 2c) strongly increased with this continuum, while species mean community N concentration (Fig. 2d) showed a weak decrease. Species’ community P concentration appeared as a strong predictor for community N:P, predominantly explaining an increasing trend (Fig. 2e). This result shows that threatened species are associated with nutrient-poor, P-impoverished habitats (14, 18), with community N:P ratios decreasing along the extinction-invasion continuum. Furthermore, our study confirms that non-invasive and invasive species elsewhere are generally adapted to nutrient-rich environments (13, 14) and have a high community P concentration and are adapted to N-limited niches (10, 14). These results indicate that nutrient conditions strongly govern species’ position along the extinction-invasion continuum and associated extinction risk and invasive success.

**Figure 2.**
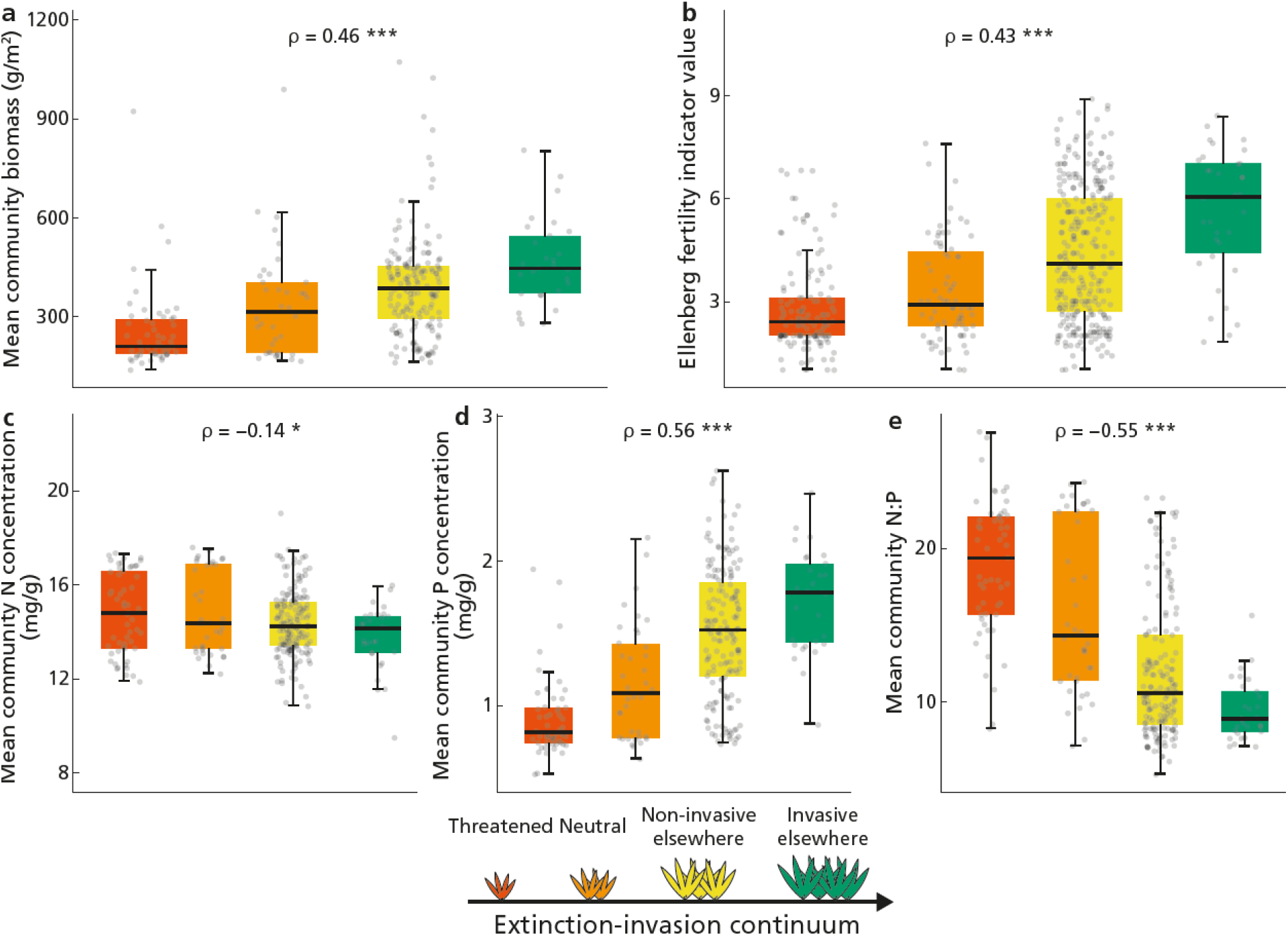
| Nutrient niche variables assessed along the extinction-invasion continuum: threatened (red), neutral (orange), non-invasive elsewhere (yellow) and invasive elsewhere (green). Nutrient niches are shown as species’ mean community biomass (**a**, n=286), Ellenberg’s soil fertility indicator value (**b**, n=525), mean community nitrogen (N) concentration (**c**, n=286), mean community phosphorus (P) concentration (**d**, n=286) and mean community N:P ratio (**e**, n=286). Boxplots display the median (center line), interquartile range (box), and whiskers extending to 1.5× the interquartile range. Outliers are shown as individual points. Spearman’s ρ is shown with its P-value (***P<0.001; *P<0.05).

Second, our results show that the proposed extinction-invasion continuum is reflected in traits related to growth rate and life history. Along with increasing a species’ extinction-invasion continuum position, species increasingly exhibited a greater maximum height (Fig. 3a) and showed a weak correlation with specific leaf area (SLA, Fig. 3b). Regarding life-history traits, our results show that naturalized and invasive species are more often annuals and biennials than perennials (Fig. 3j) and invest more in sexual reproduction, as indicated by an abundant production of small seeds (Fig. 3f, g) with a longer flowering period (Fig. 3i). Contrary to our expectations (Fig. S1), there was no correlation between a species’ position along the continuum and their belowground traits. A high root tissue density (RTD, Fig. 3d) indicates a conservative strategy, whereas a high mean root diameter (MRD, Fig. 3e) suggests a habitat favorable for mycorrhizal association, reflecting a cooperative strategy (27). A previous study showed that nutrient-poor P-limited communities are more conservative (higher RTD) and collaborative (higher MRD) than nutrient-rich N-limited communities (24). Our results show that the status of species along the extinction-invasion continuum is not reflected in these belowground traits. However, the availability of belowground traits for our species is relatively low (<30%), particularly for threatened and neutral species (<10%), stressing the need for improved sampling of these groups.

**Figure 3.**
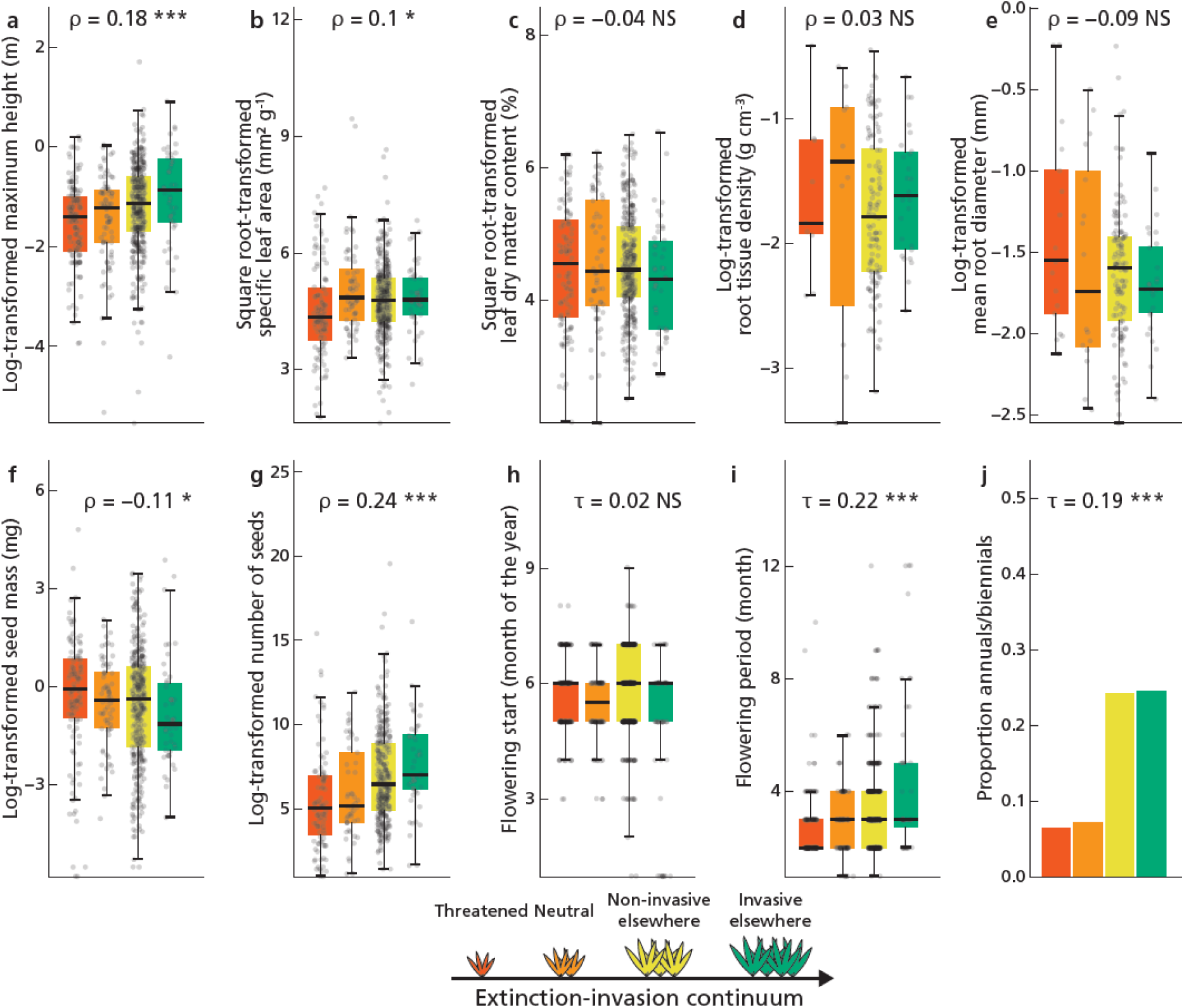
| Ten trait values assessed along the extinction-invasion continuum: threatened (red), neutral (orange), non-invasive elsewhere (yellow) and invasive elsewhere (green). Traits from left top to bottom right are maximum plant height (**a**, *n*=614), specific leaf area (SLA, **b**, *n*=548), leaf dry matter content (LDMC, **c**, *n*=535), root tissue density (RTD, **d**, *n*=166), mean root diameter (MRD, **e**, *n*=186), seed mass (**f**, *n*=562), seed number (**g**, *n*=477), flowering start (**h**, *n*=626), length of flowering period (**i**, *n*=626) and life span (proportion annuals/biennials vs perennials, **j**, *n*= 636). Boxplots display the median (center line), interquartile range (box), and whiskers extending to 1.5× the interquartile range. Outliers are shown as individual points. Spearman’s ρ is shown for continuous traits and Kendall’s T for interval or binary data with its P-value (***P<0.001; **P<0.01; *P<0.05; NS, not significant).

Third, to assess the effect of threat, naturalized, and invasive status on species’ traits and nutrient niches, we compared these groups directly (Fig. 4). Within an extended dataset of (15), we confirm that threatened species were more common in P-limited, nutrient-poor niches and exhibited distinct functional traits. They are more often annuals or biennials, with slightly lower maximum height, SLA, seed number, and a shorter flowering period (Fig. 4a). In direct comparison with invasive species (Fig. 4b), threatened species occupy distinct nutrient niches, while invasive species have relatively larger maximum height and higher SLA, produce more seeds and have a longer flowering period. While one study comparing 13 threatened and four invasive species found initial evidence for distinct nutrient niche patterns (14), our analysis provides robust support for this pattern across a broad set of European species. These results suggest that the mechanisms underlying extinction risk and invasion success in Europa are two sides of the same coin.

**Figure 4.**
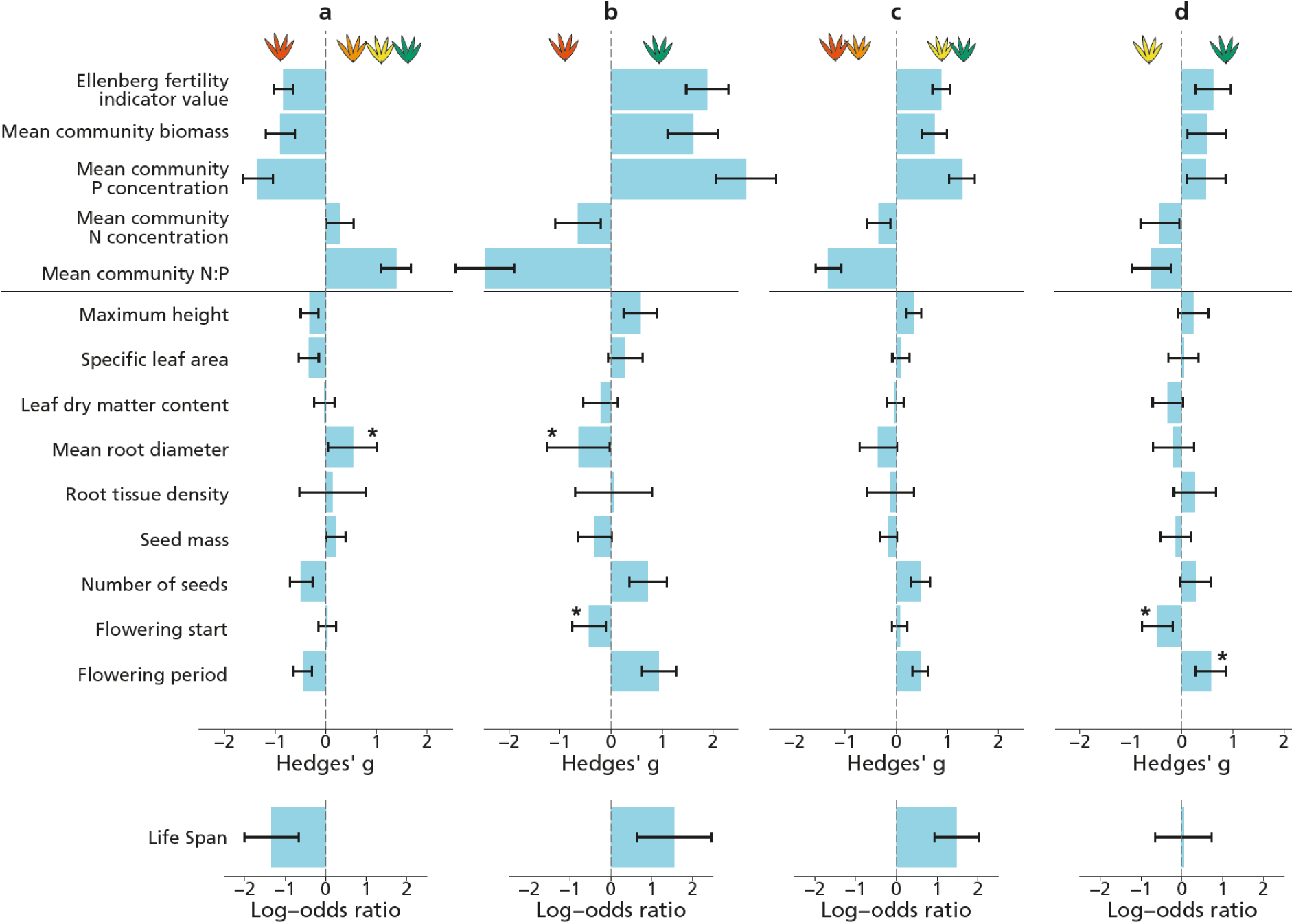
| Effect of species status on nutrient niche and trait values. Difference in traits and niches was calculated between threatened and non-threatened species (**a**), threatened and invasive species (**b**), naturalized and non-naturalized species (**c**) and naturalized non-invasive and naturalized invasive species (**d**). The traits are from top to bottom: Ellenberg’s soil fertility indicator value, species’ mean community biomass, mean community nitrogen concentration, mean community phosphorus concentration, mean community nitrogen-to-phosphorus ratio, maximum plant height, specific leaf area (SLA), leaf dry matter content (LDMC), mean root diameter (MRD), root tissue density (RTD), seed mass, seed number, flowering start, length of flowering period and life span (annuals/biennials vs perennials). Effect of species ‘status was assessed by calculating Hedges’ g for continuous and interval values and log-odds for life span. Positive values indicate higher value for the associated status indicated by plant color above the graph. Whiskers show a 95% confidence interval. Asterix next to the bar indicated that assumptions of homogeneity of variances and normality were violated and Welch’s t-test or Wilcoxon rank-sum test were non-significant (P>0.05) and results differed from Hedges’ g.

Furthermore, the distinct nutrient niches and functional traits of species that are threatened in their native range and those that naturalize elsewhere indicate that direct competition between threatened and invasive species is unlikely to be the primary driver of extinction (28) as both groups exhibit distinct trait strategies aligned with their specific nutrient requirement. Although threatened and invasive species may temporarily coexist in the same plant communities, the successful establishment of invasive species appears to depend on increased nutrient availability and N:P stoichiometry, with phosphorus fertilization being a key driver (29). When comparing species that have naturalized outside Europe with those that have not, the former are predominantly associated with nutrient-rich, nitrogen-limited environments and tend to have greater maximum height, seed production, and longer flowering periods (Fig. 4c). Traits alone cannot explain why some naturalized species become invasive while others do not; however, invasives are more likely to inhabit a nutrient-rich N-limited environment (Fig. 4d). This suggests that naturalization is potentially trait-driven, as species with greater propagule pressure have a higher chance of successful transport and introduction (7), but their invasiveness within the donor region is less likely to be explained by traits.

## Role of P in threatened or invasive status

Overall, plant traits were less effective than species’ nutrient niches in explaining their positions along the extinction–invasion continuum. This implies that we are potentially missing some key processes explaining why some species are ecological winners and others are losers. Phosphorus concentration in the community biomass was a strong predictor for species’ position along the extinction-invasion continuum, ranging from a generally low concentration for threatened species to a generally high concentration for species that are invasive elsewhere. The low community P concentration of threatened species may relate to an inherently slow growth of these species, as the relative growth rate (RGR) is strongly correlated with leaf P concentration (30) and a conservative growth strategy (15). The much higher predictive value of community P concentration for species’ position along the extinction-invasion continuum compared with specific leaf area (which also strongly correlates with RGR (31)) highlights that variation in maximum RGR is not the sole driving force and that other factors specifically linked to community P concentration also play a significant role. For example, high leaf P concentration is attributed to abundant ribosomal RNA and phospholipids (32), which allow rapid protein synthesis and turnover (33). Therefore, a high leaf P concentration likely enhances an invasive species’ plasticity (34) and may explain why these successful species are more often found in disturbed anthropogenic ecosystems (10). Furthermore, invasive species can accumulate P beyond their requirements (35); yet, it remains unclear whether this also occurs in species within their native range. Finally, P availability also plays an important role in the investment in sexual reproduction, as seeds are relatively P-rich (36). Interspecific variation in this investment appears important for a species’ positioning along the extinction-invasion continuum and merits further study.

## Join the locals

Our results show that species naturalized outside Europe are adapted to nutrient-rich and N-limited niches. This raises the question of whether non-European species naturalized within Europe exhibit similar preferences and whether they may pose a direct threat to threatened European species. To test this, we identified 16 species in our dataset from non-European origin that occur in Europe (Table S1). We analyzed these species for their nutrient niches and concluded that non-European species naturalized in Europe exhibited a more fertile niche (Fig. 5b) and a lower community N:P ratio (Fig. 5c) than threatened European species. Our results seem to mirror patterns from the Brazilian cerrado, where invasive species persisted in plant communities with low N:P ratio and occupied nutrient niches distinct from threatened species (14), indicating similar underlying adaptations to nutrient availability that facilitate non-native invasion. Despite these niche differences, we observed no significant difference in biomass production between communities dominated by non-European naturalized species and those with threatened natives (Fig. 5a). The relatively low biomass response in communities invaded by non-European species with high Ellenberg soil fertility values suggests that nutrient enrichment is still in its early stages, and that biomass accumulation is experiencing a slight lag (37). Non-European species currently exhibit relatively low abundance within these invaded plant communities (less than 10% cover), but their presence may increase over time.

**Figure 5.**
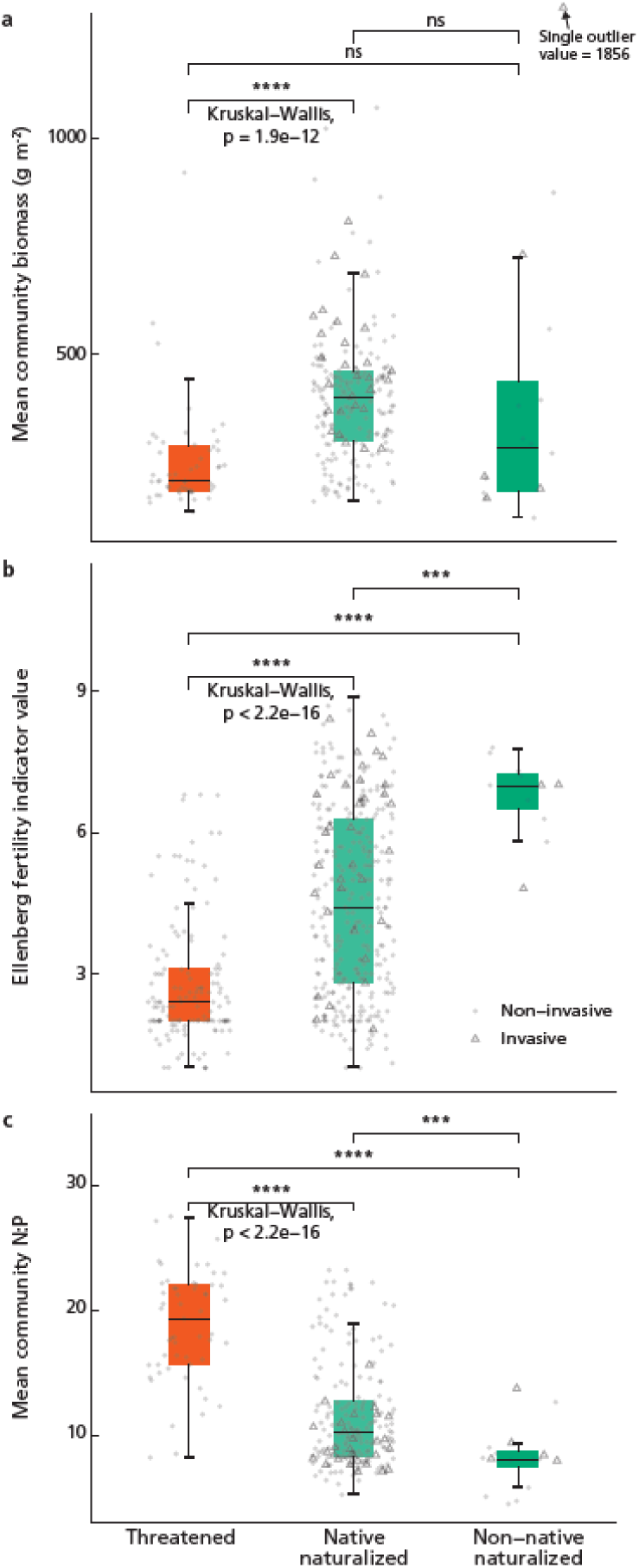
| Comparison of three nutrient niche variables between threatened species, naturalized species native to Europe and 16 naturalized species not native to Europe. Species’ nutrient niches are shown as their mean community biomass (**a**, n= 55, 231 and 16), Ellenberg’s soil fertility indicator value (**b**, n= 136, 390 and 11) and mean community nitrogen (N) to phosphorus (P) ratio (**c**, n= 55, 231 and 16). Boxplots display the median (center line), interquartile range (box), and whiskers extending to 1.5× the interquartile range. Outliers are shown as individual points. We assessed differences between groups using a Kruskal– Wallis test, followed by pairwise Wilcoxon rank-sum tests with Bonferroni adjustment for multiple comparisons (**** P < 0.0001; *** P < 0.001; NS, not significant).

Native patterns of species abundance in the Netherlands mirror global success along the extinction–invasion continuum (Fig. S2), supporting the idea that species abundant in their native range are more likely to become successful invaders elsewhere (38). Successful species that have either naturalized inside or outside Europe show a relatively similar suite of traits: investment in rapid growth and seed dispersal (11, 39). As nutrient niches and traits are relatively similar between successful native and non-native species, it indicates that demarcating between non-native and native species is poorly predictive as plant success is less likely a function of geography and more likely a function of a suite of traits that would enable this success in anthropogenic nutrient-rich environments (40): our European herbaceous species seem to join the locals in their nutrient-rich habitats (41). Nevertheless, geography may still appear to play a role in the impact of naturalized species within these environments, as invasive species can show more dominant performance when introduced in a non-native range compared with their native range (42). This improved performance of invasive species may result from escape from native competitors, herbivores and soil pathogens (43), potentially providing short-term economic advantage or its introduction of novel weapons (such as chemical toxins and allelochemicals) to which native species have not adapted (44).

## Environmental pressure, phylogeny and scope

To further assess the role of nutrient niches in the extinction–invasion continuum, we examined whether other environmental pressures contribute to the observed variation. Using ordinal regression with Ellenberg soil fertility and other Ellenberg indicator values as predictors (Table S2), we found that soil fertility remained the strongest predictor of a species’ position along the continuum. Specifically, species associated with higher soil fertility were more likely to be invasive and less likely to be threatened. Furthermore, species preferring wetter or more alkaline conditions were less likely to be invasive, while those favoring higher light levels showed a greater likelihood of being invasive. The influence of these environmental indicators on the success of native species has already been well established (45, 46), but our results suggest they also promote naturalization and invasion. Next, we investigated whether phylogenetic relatedness influenced species’ positions along the extinction-invasion continuum. We found a moderate phylogenetic signal (Pagel’s λ = 0.58), indicating that closely related species tend to occupy similar positions along the continuum. Non-invasive elsewhere status was phylogenetically widespread across Poaceae, Brassicaceae, Fabaceae, Plantaginaceae and Asteraceae, whereas threatened status is broadly distributed across Cyperaceae and Orchidaceae (Fig. S3, Table S3). A previous study (47) reported that naturalized species are often a phylogenetically random subset of their source species pools, particularly in human-influenced habitats. Our results indicate that although closely related species tended to share naturalization status, the effect was only moderate, suggesting that naturalization is likely shaped by a combination of ecological, evolutionary, and anthropogenic factors

Species in our dataset have mainly been introduced to temperate regions (Fig. S4), which are identified as the main hotspots of species introduction and naturalization (48). However, it remains unclear if the observed patterns also hold for Mediterranean, subtropical or tropical regions. Early studies have indicated nutrient enrichment is a prerequisite for invasive success in Mediterranean, subtropical and tropical regions (14, 29, 49). More research in these regions is critical as they are often hotspots of biodiversity (50) and support the persistence of threatened species, of which many are associated with old P-impoverished soils (51). Our dataset comprises relatively young soils at higher latitudes where glaciations have occurred quite recently, bringing P-rich sediments to the surface. In these regions, P limitation occurs mainly on soils with specific biogeochemical conditions such as areas where calcareous groundwater comes to the surface, or in areas where P impoverishment has taken place by hay harvesting in absence of fertilization Many species on older P-impoverished soils, such as in Australia, South Africa, and Brazil, show limited seed dispersal and strong fire-responsive strategies (52) as well as diverse nutrient-acquisition strategies (53), potentially explaining the relatively low numbers of naturalized species in these soils (although examples of naturalized plant species exist (54)). In our dataset, there are several species that persist under low-P conditions (N:P>16) that have naturalized outside Europe (n=31, 16.1%). We analyzed how the extent of naturalization (abundance on naturalization lists) related to species’ nutrient niche (Fig. S5). The degree of naturalization was negatively correlated with species’ N:P ratios and positively correlated with Ellenberg’s soil fertility indicator values. Additionally, P-limited naturalized species were not invasive and exhibited a low naturalization extent. This suggests that naturalized P-limited species originating from Europe pose a limited threat to non-European P-limited species. Investigating whether naturalized species occupy similar nutrient niches in their naturalized and native ranges represents an important avenue for further study. Furthermore, the suggestion that invasive species are unable to outcompete European threatened species requires further experimental validation by testing their performance and competition under varying nutrient levels.

## Mitigating and preventing nutrient enrichment

Currently, the nutrient dynamics are globally changing due to N and P inputs (55). Naturalized species, either invasive or non-invasive, are disproportionately advantaged in such a nutrient-enriched world (56). Previous studies (18, 57) urged limiting P fertilization to prevent species loss, especially as P is a finite resource (58). We contribute to this concern by indicating that, within the context of the extinction-invasion continuum, induced nutrient-enriched and N-limited environments potentially increase the chance of species invasion. Phosphorus fertilization induces a shift from P to N limitation (55) whereas N deposition can increase overall nutrient availability and productivity (59) and, in light of this study, both fertilization processes are a potential driver of species naturalization as well as species extinction. Furthermore, an increase in naturalized species is associated with a higher likelihood of invasive species establishment (48), also visible in our data (Fig. S4), and combined with nutrient enrichment, can potentially lead to serious ecological changes. However, our results suggest that within the context of Europe, invasive species do not pose a direct threat to threatened species and species replacement occurs through local nutrient regime shifts. Therefore, mitigating and preventing nutrient enrichment, especially phosphorus, serves a dual purpose in nature conservation by protecting existing nutrient habitats for threatened species and simultaneously limiting opportunities for invasive species to establish.

## Acknowledgments

We thank Ton Markus for improving the figures. We thank Lenze Hofstee for providing us the opportunity to stay in the red brick house in Kempenbroek, where this idea was conceived and further refined.

## Author contributions

DS conceived the idea. DS, HO and HB developed the conceptual models underlying this study. DS designed the methodology and analyzed the data. DS and HO made an outline for the paper, DS wrote the original draft and HO helped significantly. HO, HB, JS, HL, MW and PB were involved in multiple critical discussions on drafts. All authors critically reviewed the drafts and provided comments. HO, HB, JS, and MW took on a supervising role.

## Data and materials availability

All data and codes supporting the results presented in this manuscript are on a OSF repository and will become available after publication.

## Supplementary Materials

Materials and Methods

Figs. S1 to S6

Tables S1 to S5

## Supplementary Materials

## Materials and Methods

### Sampling procedure

The dataset used in this study comprises data from 1038 European sites of herbaceous vascular plant communities (15, 24) that were collected in multiple countries: the Netherlands (n=617), Belgium (n=20), Poland (n=169), Russia (n=83), Germany (n=90), UK (n=16), Sweden (n=16), Belarus (n=10), and Iceland (n=17). The sites encompass a variety of conditions, ranging from dry to wet environments, and include herbaceous grasslands, fens, bogs, marshes, reedbeds, and dune vegetation. The sampling was carried out within plots of varying sizes (0.06–26 m²), which were chosen to best represent the species composition of each site.

Species composition and cover were recorded, and a 20x20 cm subsample was taken during the peak growing season (June–August). The sampling year of the plots varied across the dataset and occurred between 1990 and 2020. The aboveground living biomass of plants in this subsample was harvested, and their dry weight was determined (g m^-2^). The concentration of nitrogen (N), phosphorus (P), and potassium (K) in the aboveground plant material was then analyzed, with N measured using an NCS analyzer, and P and K quantified by ICP-OES.

### Nutrient niche and traits

From all available plots (n=1038), we omitted plots that were outside Europe (Russian Siberia, n=83), that were woody (cover woody species > 50%, n=37), or that showed signs of K-limitation (60) (N:K > 2.1 and K:P < 3.4; n=81). Within the remaining 837 plots, we recorded 682 unique species. We retrieved traits related to growth and life history by building on previous studies (15, 24) and complemented new species trait data on their original and standardized name using trait databases LEDA (61), BIOBASE (62) and GIFT (63) (see repository for data availability and Table S4). Several traits were log-transformed or square-root transformed to improve the interpretability and visualization of their distributions. We used community aboveground biomass as a proxy for primary productivity and hence overall nutrient availability (64, 65), and community aboveground biomass N:P:K as a proxy for relative nutrient availability (66, 67). We complemented species with Ellenberg’s soil fertility indicator values (23) by matching our species with this dataset for species that had a European Ellenberg nutrient indicator value available (79% of species). For species that were considered synonyms, we used the value calculated for the original species name. Furthermore, we calculated species’ mean nutrient niche positions based on community aboveground biomass and nutrient concentrations for species that occurred in at least ten plots. Sixteen non-native species that have naturalized in Europe were only marginally found in the dataset (between one and eleven plots), and for these we computed the mean regardless of any minimum threshold. We excluded all woody species from our analysis.

### Extinction-invasion continuum

We approached the extinction-invasion continuum by assigning four status (1= threatened, 2= neutral, 3= non-invasive elsewhere, 4= invasive elsewhere) to the species based on its presence in several databases (see Fig. S6 for an overview). Extinct species were not assessed due to the lack of observations. Non-native species that have naturalized from outside Europe (n=16) were excluded from the continuum by using a Dutch flora database (68) and we manually checked for any remaining species (Table S1). Next, we determined if species in our dataset were naturalized outside Europe by checking their presence in the global naturalized flora database (GLONAF) (25) for all non-European countries as defined by the database. We also recorded the naturalization extent of that species, that is the number of lists the species was found (similar to (69)). Continuing, these naturalized species were labeled as invasive elsewhere if they were present in a global invasive database (GIDS) (26) and were labeled as non-invasive elsewhere if they were absent. For the remaining species, the threatened status was based on the presence of this species in a combined red list of the eight countries (the Netherlands, Belgium, Poland, Germany, UK, Sweden, Belarus and Iceland) in which our communities were sampled (similar to (18), see the repository for data availability). Species were assigned as threatened if they were identified as ‘critically endangered’, ‘endangered’ or ‘vulnerable’. Finally, we determined a species as neutral if it was neither determined as naturalized (either invasive or non-invasive) or threatened. Species names in all datasets were standardized according to World Flora Online backbone using the Taxonomic name resolution service (70). We compared the nutrient niches of threatened, native, and non-native naturalized species using the Kruskal-Wallis test (for overall differences across all three groups) followed by post hoc pairwise Wilcoxon rank-sum tests with Bonferroni correction. We examined the effect of species’ position within the extinction-invasion continuum on nutrient niche indicators (Ellenberg’s soil fertility indicator value and mean community aboveground biomass, N concentration, P concentration and N to P ratio) and continuous traits by calculating Spearman’s ρ. We used Kendall’s T when trait data were interval or binary. Species’ life span was analyzed using binary values (annuals and biennials as 1 and perennials as 0), but is depicted in this study as proportions.

### Effect of species’ status on traits and niches

Next, we investigated the effect of threatened, naturalized and invasive status on traits and nutrient niches by comparing several groups directly using a group-wise comparison. For this, we adopted a method similar to (15), where we calculated the relative difference between traits and niches quantified by an effect size and a 95% confidence interval. For interval and continuous traits and all niches, Hedges’ g was used as the effect size measure, as it corrects for bias due to unequal sample sizes between groups. For species’ lifespan (annuals/biennials vs perennials), we calculated log-odds ratio. We assessed assumptions of normality and homogeneity of variances. Interval traits (flowering period, start of flowering) are discrete and measured in equal intervals and therefore normality testing is not appropriate. When assumptions of homogeneity of variances were violated, we checked our results by applying Welch’s t-test. When normality assumptions were violated, we checked our results by applying a Wilcoxon rank-sum test to compare groups. Significant differences were reported in the main figure.

### Ellenberg values and abundance in the Netherlands

We aimed to test whether species’ position on the extinction-invasion continuum can be strongly predicted by Ellenberg nutrients when we account for five other Ellenberg values using an ordinal regression model. We complemented our dataset with five other Ellenberg indicator values (temperature, light, salinity, moisture, and reaction) by matching species in our dataset to a European Ellenberg database (23), using only species for which values were available. For species that were considered synonyms, we used the value calculated for the original species name. Next, we ran an ordinal regression model (71) in R. We included the Ellenberg indicator values as predictors and the ordinal position of species on the extinction-invasion continuum as the response variable. As salinity was heavily left-skewed in our dataset, only 8% of species had an Ellenberg salinity value above 1 and therefore this variable showed strong zero-inflation, potentially complicating model interpretation. To assess the robustness of our results, we reran the model excluding Ellenberg salinity, which yielded similar conclusions.

Next, we were interested in testing how the extinction-invasion continuum relates to species’ general abundance at home. We adopted an analysis to test such a relationship for the Netherlands. The list of Dutch flora from 2020 (68) has information on an estimation of the abundance of species within 1-km grid cells (K) for the period 1902 –1949 (KFK1930), 1975 –1987 (KFK1980), 1988 –1999 (KFK1995) and 2000 – 2019 (KFK2015). The abundance estimation is expressed as KFK, which is a value between 1-9, where 1 consists of 1-3 K’s and 9 consists of more than 10.000 K’s (see Table S5). We added this information to our dataset, with 483 (73%) species showing a KFK classification for KFK19. We analyzed the relationship between species positions along the extinction-invasion continuum and their KFK scale by calculating Kendall’s T for the four periods.

### Phylogeny

We aimed to visualize the phylogenetic structure in our dataset and test whether the four positions (threatened, neutral, non-invasive elsewhere and invasive elsewhere) along the extinction-invasion continuum show a clustered or random phylogenetic pattern. First, we added information on the family of the plants from the World Flora Online backbone. Next, we constructed a phylogenetic tree for our species using the package V.PhyloMaker2 (72). We used ‘scenario 3’ to add missing species (21%) to our tree by attaching them to the most appropriate positions based on their genus or family-level taxonomy. We excluded subspecies from our tree. Species of the genus *Rumex* were not present in the backbone phylogeny, so we manually added a genus-level node for *Rumex* under the family Polygonaceae. This resulted in a tree from 651 species. Next, we tested whether species’ position along the extinction–invasion continuum showed a phylogenetically structured pattern using Pagel’s lambda (λ) (73) from the phytools package (74). Pagel’s λ is a phylogenetic scaling parameter used to assess how well the evolutionary relationships among species explain the species position along the extinction-invasion continuum, where λ = 1 indicates a strong phylogenetic signal consistent with so-called Brownian motion evolution, and λ = 0 indicates no phylogenetic signal (random distribution across phylogeny).

---

**Figure S1.**
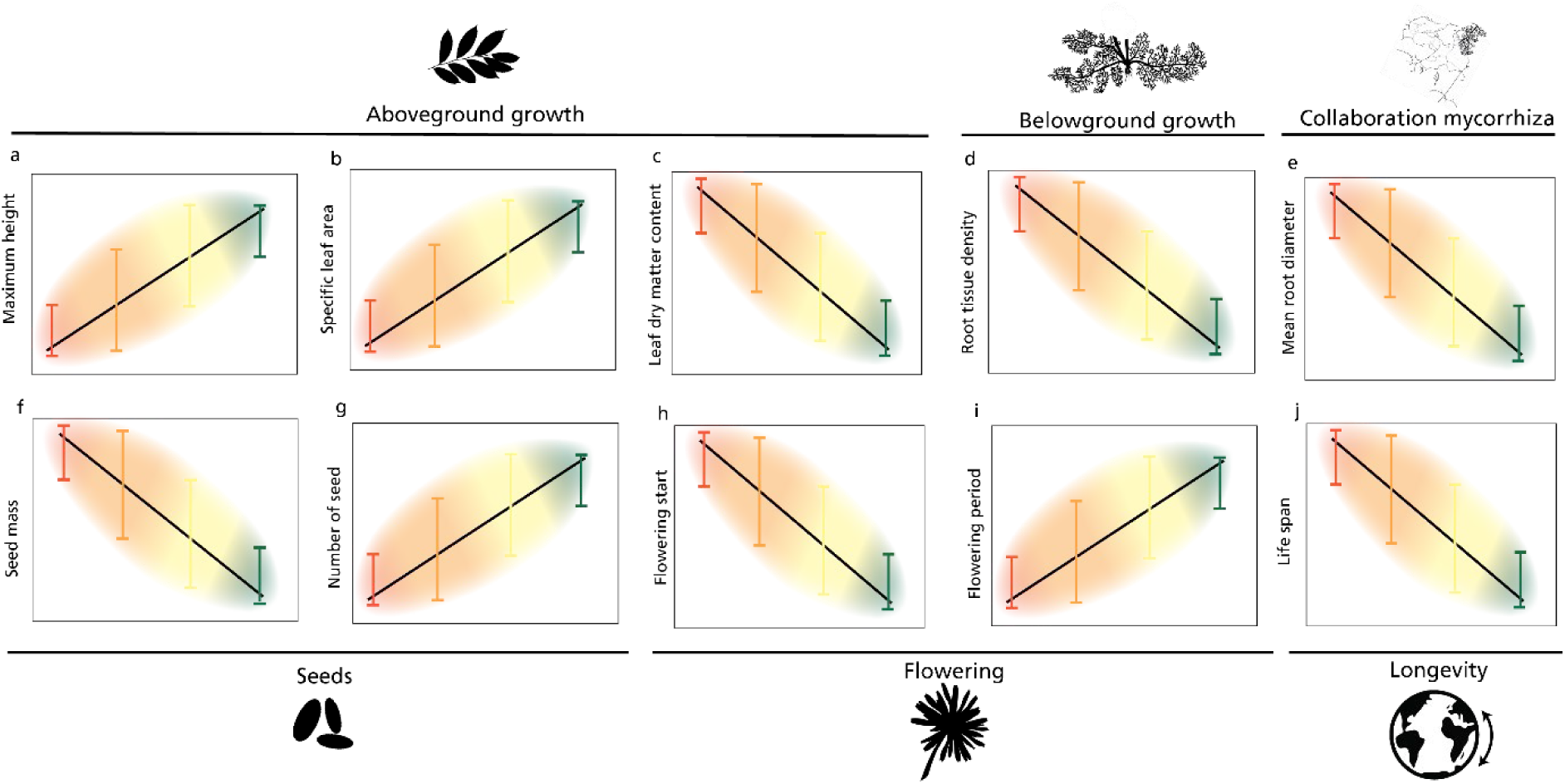
| Expected relationships between traits and species’ positions along the extinction-invasion continuum. Our general expectation is that naturalized invasive (green) and non-invasive species at the end of the extinction-invasion continuum invest more in fast growth and return and are more focused on life history traits allowing a quick dispersal strategy compared to neutral (yellow) or threatened species (red). Considering aboveground and belowground growth traits along the extinction-invasion continuum, we expect maximum plant height (**a**) and specific leaf area (**b**) to increase, indicative of fast biomass accumulation. Leaf dry matter content (**c**) and root tissue density (**d**) should therefore decrease as it is associated with investment in maximizing the longevity of tissue. Furthermore, as we expect that threatened species are more common in phosphorus-limited conditions, we expect more collaboration with mycorrhiza depicted by an increase in species’ root diameter (**e**), providing habitat for mycorrhizal association. Next, we anticipate that species will enhance their reproductive efforts towards the end of the extinction-invasion continuum visible in species’ seed investment and sexual reproduction. We expect that species along the extinction-invasion continuum have smaller seed mass (**f**) produced in larger numbers (**g**), enabling a higher chance to invade and colonize new areas. Additionally, as the extinction-invasion continuum increases, species will exhibit a lower flowering start (**h**), indicating an earlier start of flowering, and a higher flowering period (**i**), reflecting a longer period in which the species flowers. Finally, as invasive and naturalized species are more associated with annuals or biennials, showing lower life longevity, we expect that the lifespan decreases along the continuum (**j**). Silhouette images were obtained from Phylopic.

**Figure S2.**
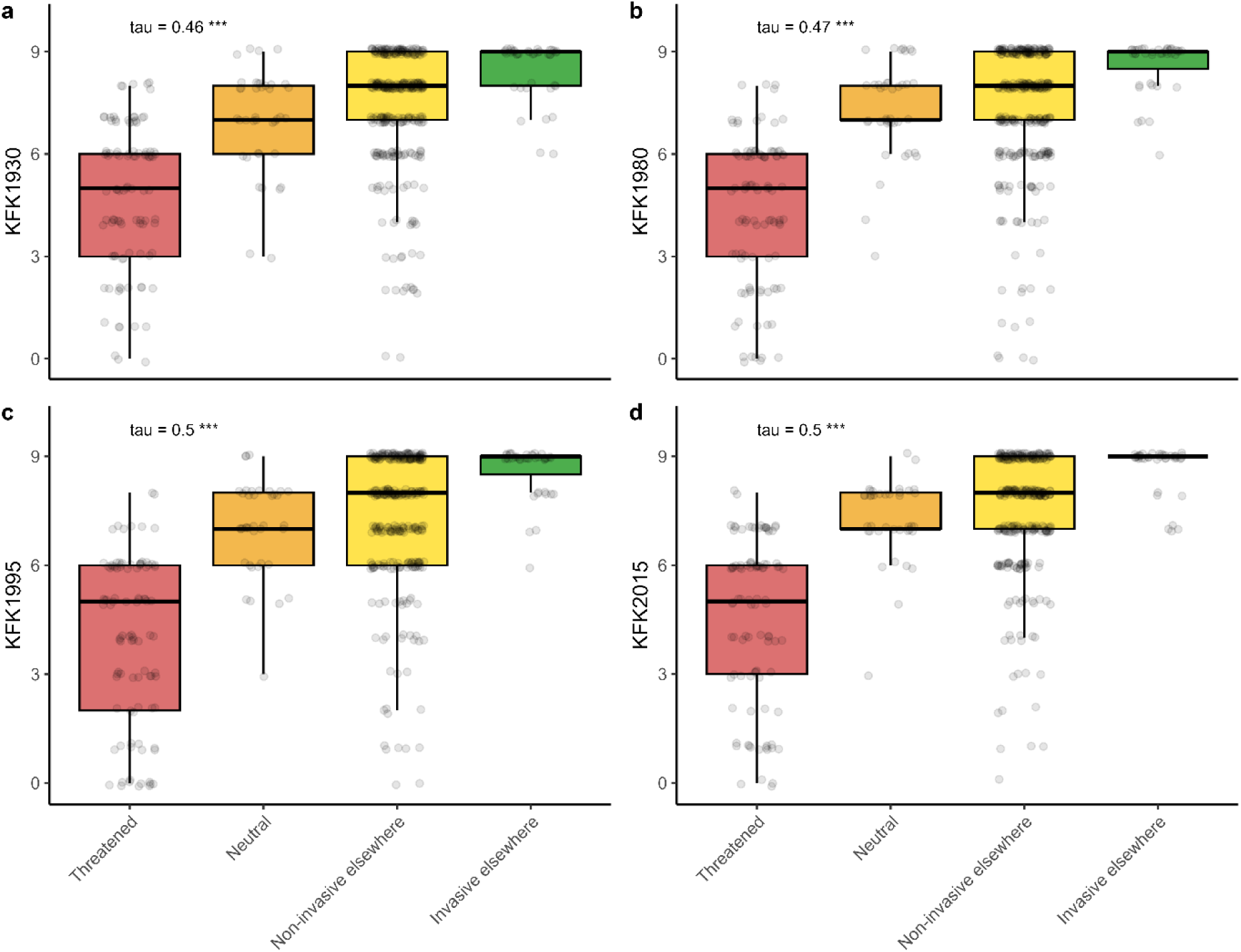
| Species positions along the extinction–invasion continuum in relation to their abundance across 1-km grid cells in the Netherlands. We plotted species positions along the extinction-invasion continuum related to Dutch estimate scale of abundance in 1-km grid cells for the periods 1902 –1949 (KFK1930, **a**), 1975 –1987 (KFK1980, **b**), 1988 –1999 (KFK1995, **c**) and 2000 – 2019 (KFK2015, **d**). The scale of the y-axis is from 0-9, see Table S5 for estimated abundances. Boxplots display the median (center line), interquartile range (box), and whiskers extending to 1.5× the interquartile range. Outliers are shown as individual points. Kendall’s T is provided on the left top with its P-value (***P<0.001).

**Figure S3.**
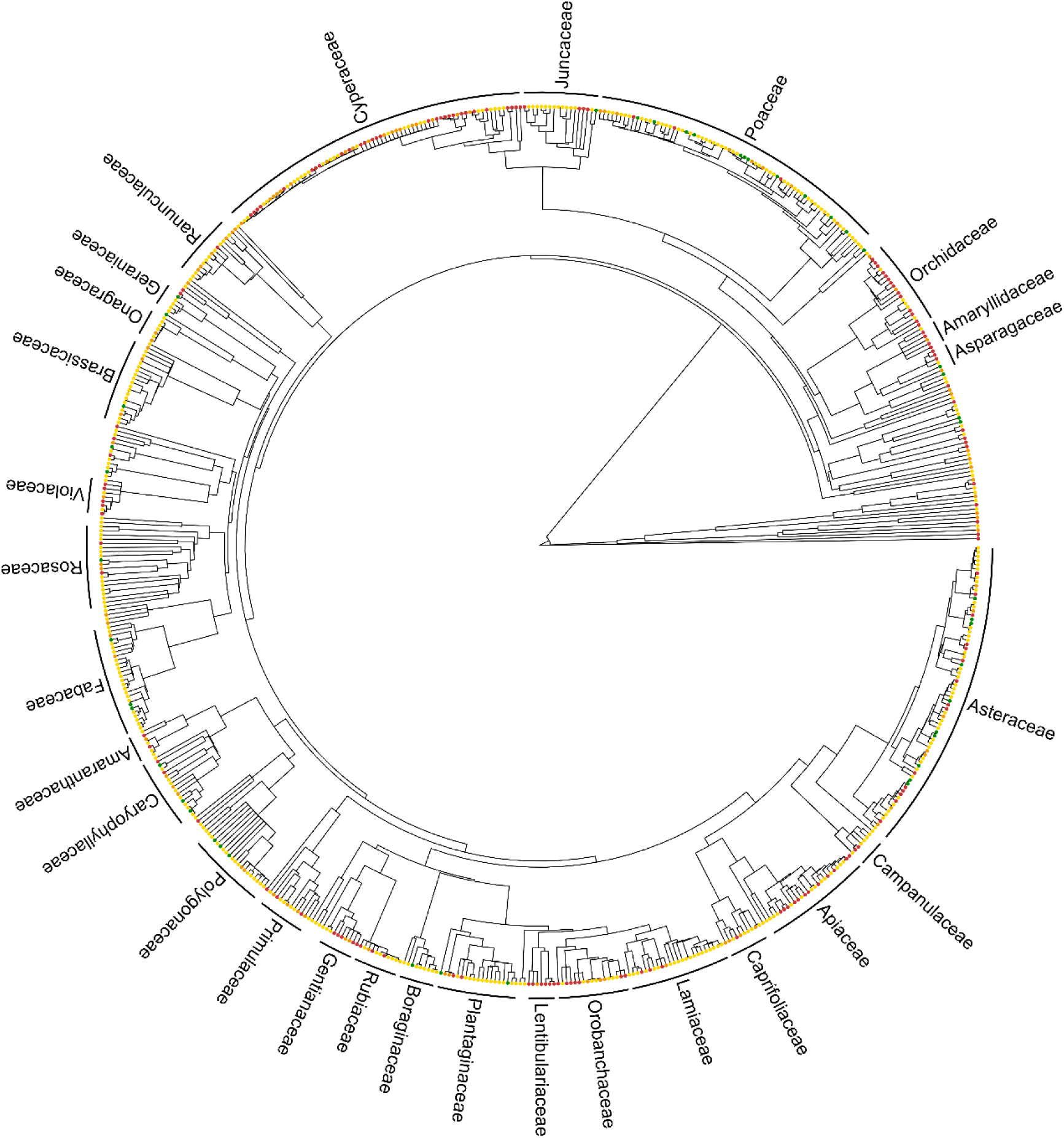
| Phylogeny of 651 species native to Europe coloured by tip. Tip colors representing species’ positions along the extinction–invasion continuum

**Figure S4.**
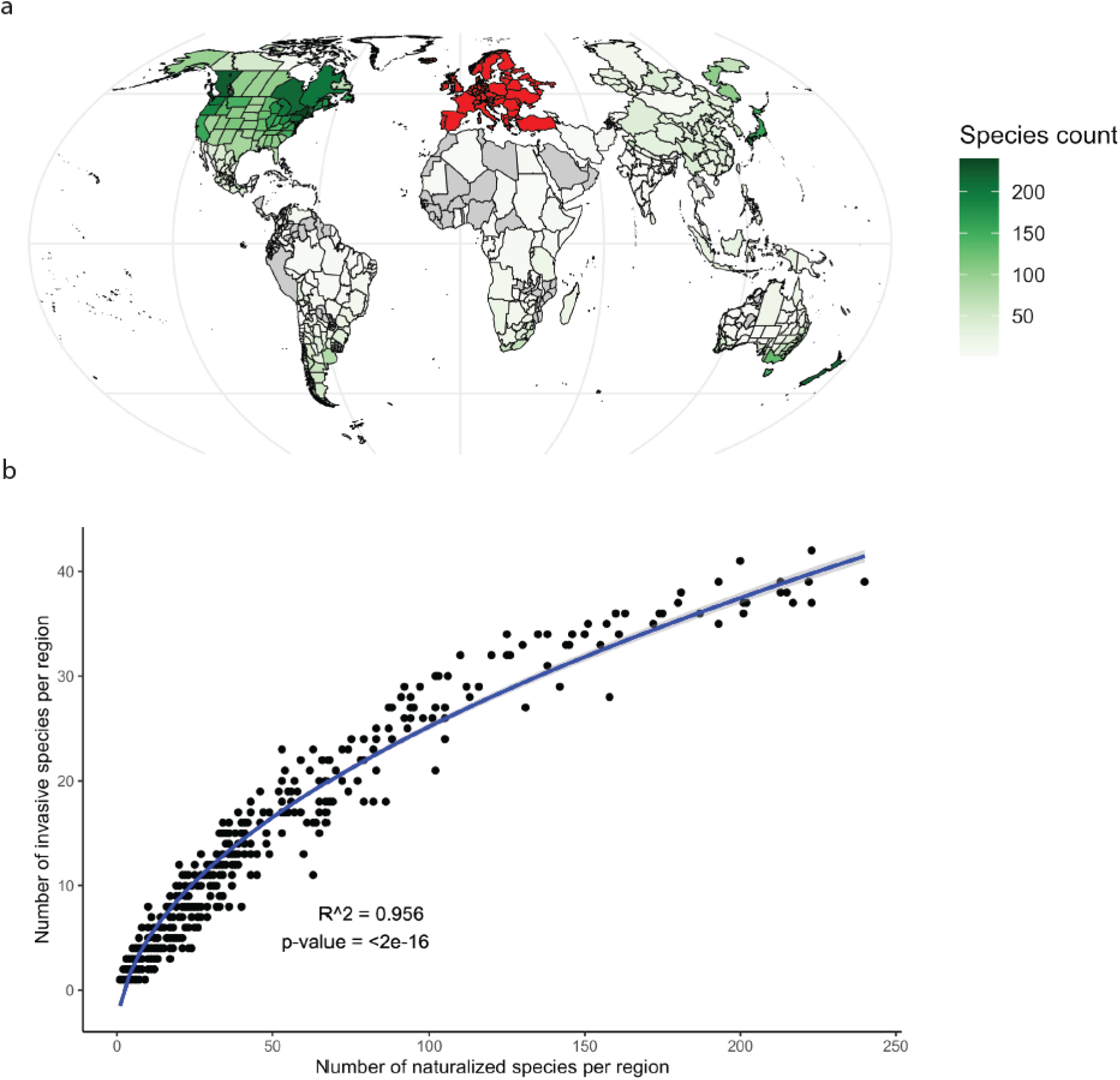
| Distribution of naturalized and invasive species from our dataset. In **a**, we show the worldwide distribution of the number of species within our dataset that have naturalized outside Europe. Regions in red were considered European and are classified as the native region. Grey areas did not have any naturalized species. Several areas were not included in GLONAF regions and are therefore missing in this map. In **b**, the number of naturalized species from our dataset per region outside Europe (x-axis) was compared with the number of invasive species from our dataset in that region (y-axis). Linear regression with a square root transformation on the predictor (y = -4.45 + 2.96*√x) is significant (p<0.001) with R^2^=0.96.

**Figure S5.**
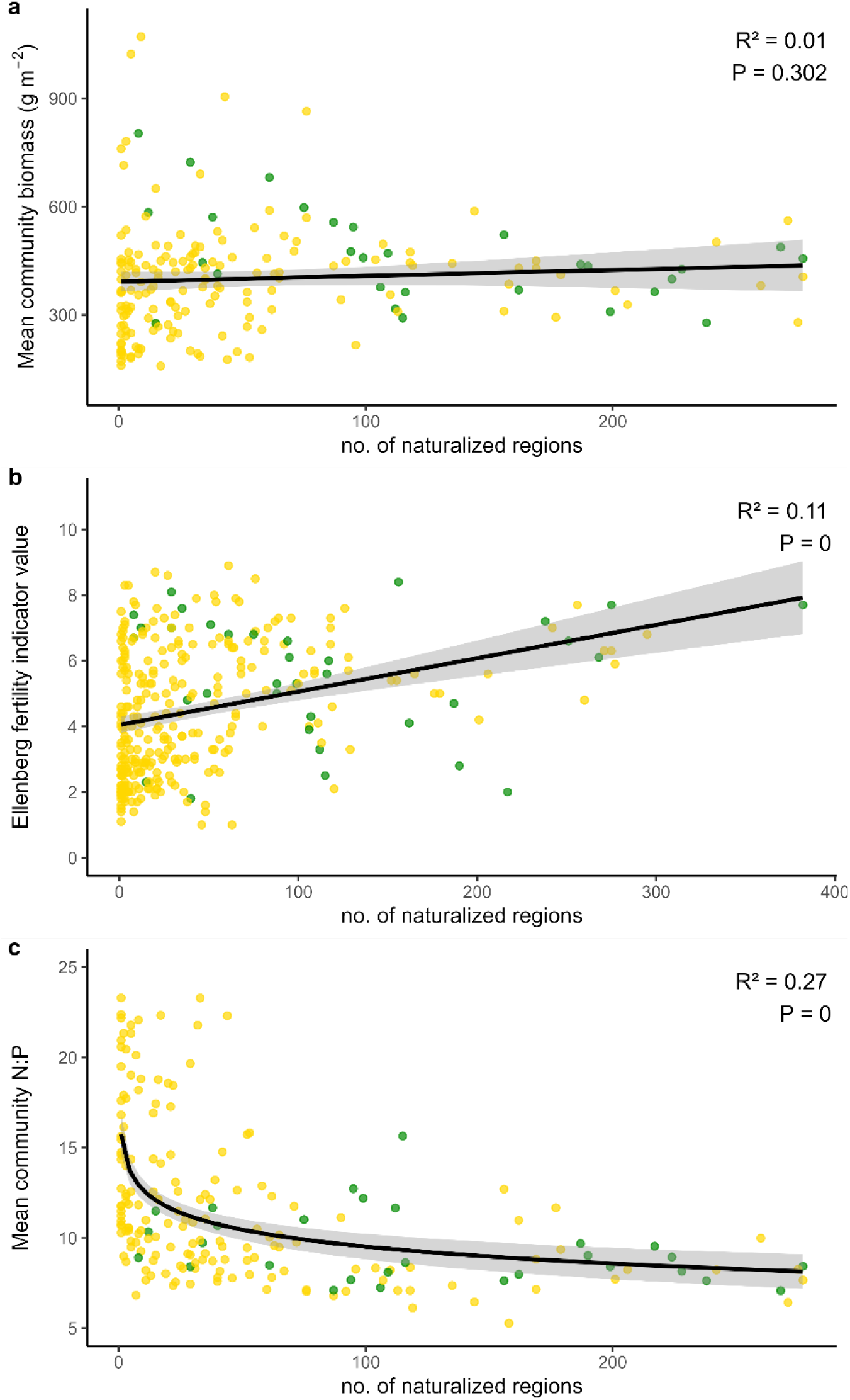
| Relationship between naturalization extent and nutrient niches. We used the count of presence per species in a region’s naturalized lists (naturalization extent) and compared it along species’ mean community biomass **(a)**, Ellenberg’s soil fertility indicator value **(b)** and species’ mean community nitrogen to phosphorus ratio (N:P) **(c).** In panels **(a)** and **(b)**, we fitted a linear model, and in **(c)**, we fitted a logistic model. R-squared and P of the models are provided in the top right-hand corner.

**Figure S6.**
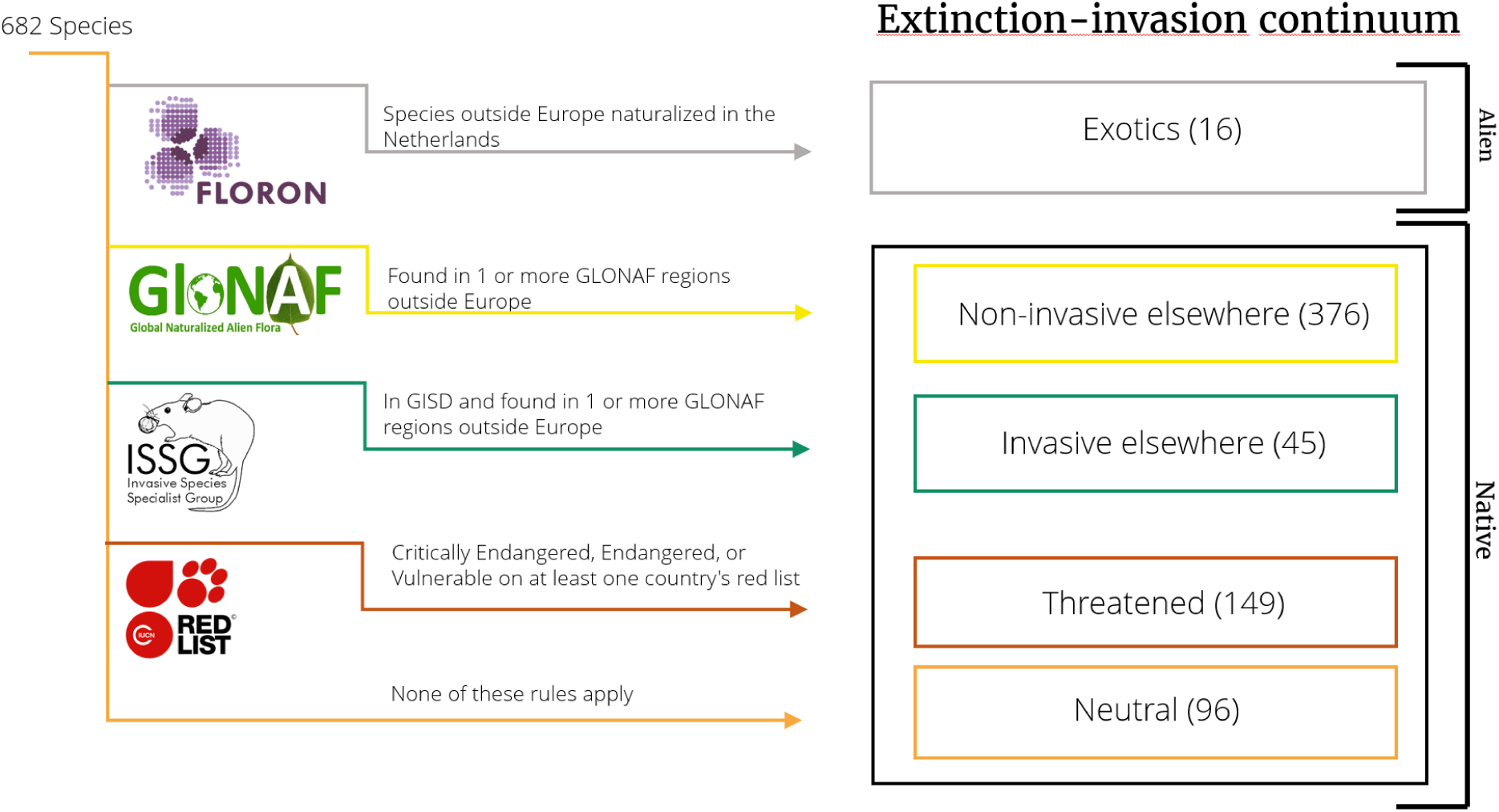
| Methodology determining species’ position along the extinction-invasion continuum. We used a partitioning structure to assign 682 species to five status. We first assigned 16 species as naturalized in Europe using the plant species database of *Floristisch Onderzoek Nederland* (FLORON) and then checked manually for any remaining species. These were outside the scope of the study. Second, 411 species were found outside Europe using the Global Naturalized Flora database (GLONAF). Of these 411 species, 45 invasive species were found in the Global Invasive Species Database (GISD) and 376 non-invasive species were not. Third, we assigned 149 species to a threatened status using the IUCN country red lists of our sites. Finally, the remaining 96 species were assessed as neutral species, as they were neither naturalized outside Europe nor defined as threatened.

**Table S1.** | Sixteen species in our dataset were identified as naturalized from outside Europe and therefore considered non-native.

| Standardized name | Origin | Invasive/non-invasive | Observed in plots (n) |
| --- | --- | --- | --- |
| **Acorus calamus** | South and East Asia | 0 | 4 |
| **Bidens frondosa** | North America | 0 | 2 |
| **Calamagrostis canadensis** | North America | 0 | 1 |
| **Crassula helmsii** | Australia and New Zealand | 1 | 2 |
| **Epilobium ciliatum** | North America | 0 | 6 |
| **Erigeron annuus** | North America | 0 | 4 |
| **Erigeron canadensis** | North America | 0 | 10 |
| **Erigeron floribundus** | South America | 1 | 1 |
| **Erigeron sumatrensis** | South America | 0 | 1 |
| **Impatiens glandulifera** | India | 1 | 1 |
| **Juncus tenuis** | North America | 1 | 2 |
| **Matricaria discoidea** | North America | 0 | 1 |
| **Rumex salicifolius** | North America | 0 | 2 |
| **Senecio inaequidens** | South Africa | 1 | 11 |
| **Solidago canadensis** | North America | 0 | 1 |
| **Solidago gigantea** | North America | 0 | 3 |

**Table S2.** | Ordinal regression results for predictors on the extinction-invasion continuum. Summary of parameter estimates, standard errors, test statistics, p-values, odds ratios, and significance levels (***P<0.001; **P<0.01; *P<0.05; NS, not significant) for ordinal regression predicting species position on the extinction-invasion continuum using Ellenberg indicator values.

| Predictor | Estimate | Std. Error | z value | Pr(>\|z\|) | Odds ratio | Significance |
| --- | --- | --- | --- | --- | --- | --- |
| Ellenberg light | 0.35 | 0.13 | 2.82 | 0.00 | 1.42 | ** |
| Ellenberg temperature | 0.05 | 0.11 | 0.32 | 0.69 | 1.05 | NS |
| Ellenberg salinity | -0.28 | 0.12 | -2.29 | 0.02 | 0.75 | * |
| Ellenberg moisture | -0.33 | 0.06 | -5.55 | 0.00 | 0.72 | *** |
| Ellenberg nutrients | 0.82 | 0.10 | 8.14 | 0.00 | 2.27 | *** |
| Ellenberg reaction | -0.31 | 0.07 | -4.31 | 0.00 | 0.74 | *** |

**Table S3.** | Extinction–invasion status distribution among plant families with >10 species in the phylogenetic analysis. Percentages are provided for species that were included in the phylogenetic analysis by family.

| Family | N species | Threatened (%) | Neutral (%) | Non-invasive elsewhere (%) | Invasive elsewhere (%) |
| --- | --- | --- | --- | --- | --- |
| **Total** | 651 | 22.3 | 14.1 | 56.7 | 6.9 |
| **Cyperaceae** | 74 | 39.2 | 25.7 | 35.1 | 0 |
| **Asteraceae** | 73 | 20.5 | 9.6 | 56.2 | 13.7 |
| **Poaceae** | 71 | 4.2 | 11.3 | 69.0 | 15.5 |
| **Apiaceae** | 26 | 46.2 | 0.0 | 53.8 | 0.0 |
| **Fabaceae** | 26 | 0.0 | 7.7 | 80.8 | 11.5 |
| **Lamiaceae** | 24 | 12.5 | 4.2 | 83.3 | 0.0 |
| **Orchidaceae** | 21 | 66.7 | 9.5 | 23.8 | 0.0 |
| **Brassicaceae** | 20 | 0.0 | 15.0 | 80.0 | 5.0 |
| **Rosaceae** | 20 | 10.0 | 10.0 | 75.0 | 5.0 |
| **Plantaginaceae** | 19 | 10.5 | 10.5 | 68.4 | 10.5 |
| **Polygonaceae** | 19 | 0.0 | 10.5 | 73.7 | 15.8 |
| **Juncaceae** | 18 | 22.2 | 0.0 | 72.2 | 5.6 |
| **Caryophyllaceae** | 17 | 11.8 | 17.6 | 52.9 | 17.6 |
| **Orobanchaceae** | 17 | 41.2 | 17.6 | 41.2 | 0.0 |
| **Ranunculaceae** | 15 | 6.7 | 33.3 | 60.0 | 0.0 |
| **Primulaceae** | 12 | 16.7 | 33.3 | 50.0 | 0.0 |
| **Rubiaceae** | 11 | 27.3 | 9.1 | 63.6 | 0.0 |
| **Caprifoliaceae** | 10 | 10.0 | 40.0 | 50.0 | 0.0 |

**Table S4.** | List of aboveground and reproductive plant traits adapted from Fujita et al. (15) **and Scheifes et al.** (24)**n and supplemented.** Ten above- and belowground traits with scale, unit, percentage of species with traits and their source are provided. Abbreviation after trait in brackets is provided for in-text and figure references. The traits were retrieved for 666 unique herbaceous species.

| Trait | Scale | Unit | % Species | Source |
| --- | --- | --- | --- | --- |
| Maximum height | Continuous | m | 92 | (61, 62) |
| Specific leaf area (SLA) | Continuous | mm<sup>2</sup> g<sup>-1</sup> | 82 | (62) |
| Leaf dry matter content (LDMC) | Continuous | % | 80 | (62) |
| Root tissue density (RTD) | Continuous | g cm<sup>-3</sup> | 25 | (75) |
| Mean root diameter (MRD) | Continuous | mm | 28 | (75) |
| Seed mass | Continuous | mg per seed | 85 | (62, 76, 77) |
| Number of seeds | Continuous | number per shoot | 72 | (62) |
| Starting month of flowering | Interval | month (Jan:1 – Aug:8) | 94 | (61, 76) |
| Duration of flowering period | Interval | month | 94 | (61, 76) |
| Lifespan | Binary | 1: annual or biennial, 0: perennial | 95 | (61-63, 76) |

**Table S5.** | estimation of abundance of species within 1-km grid cell in a 1-9 scale.

| KFK | Number of 1-km grid cells |
| --- | --- |
| 0 | 0 |
| 1 | 1-3 |
| 2 | 4-10 |
| 3 | 11-30 |
| 4 | 31-100 |
| 5 | 101-300 |
| 6 | 301-1000 |
| 7 | 1001-3000 |
| 8 | 3001-10.000 |
| 9 | >10.000 |

